# Challenge data set for macromolecular multi-microcrystallography

**DOI:** 10.1101/394965

**Authors:** James M. Holton

**Affiliations:** Department of Biochemistry and Biophysics, University of California, San Francisco, CA 94158-2330, USA; Divison of Molecular Biophysics and Bioengineering, Lawrence Berkeley National Laboratory, Berkeley, CA 94720, USA; Stanford Synchrotron Radiation Lightsource, SLAC National Accelerator Laboratory, Menlo Park, CA 94025, USA

**Keywords:** Protein, x-ray, simulation, phasing, microcrystallography, radiation damage

## Abstract

A synthetic data set demonstrating a particularly challenging case of indexing ambiguity in the context of radiation damage was generated in order to serve as a standard benchmark and reference point for the ongoing development of new methods and new approaches to solving this problem. Of the 100 short wedges of data only the first 71 are currently necessary to solve the structure by “cheating”, or using the correct reference structure as a guide. The total wall-clock time and number of wedges required to solve the structure without cheating is proposed as a metric for the efficacy and efficiency of a given multi-crystal automation pipeline.

**Synopsis:** A synthetic dataset demonstrating the challenges of combining multiple data sets with indexing ambiguity in the context of heavy radiation damage in multi-crystal macromolecular crystallography was generated and described, and the problems encountered using contemporary data processing programs were summarized.

## 1. Introduction

Data sets that challenge the capabilities of modern structure solution procedures, algorithms and software are difficult for developers to obtain for a very simple reason: as soon as a solution is reached the data set is no longer considered challenging. Data sets that are recalcitrant to current approaches are also not available in public databases such as the Protein Data Bank (Berman *et al.*, 2002) or image repositories (Grabowski *et al.*, 2016, Morin *et al.*, 2013) that only contain data used for solved structures. When testing the limits of software it is generally much more useful to know ahead of time what the correct result will be. This enables detection and optimization of partially-successful solutions at every point in the process, even if downstream procedures fail.

There is a fundamental limit to how small a protein crystal can be and still yield a complete dataset (Holton & Frankel, 2010), so as beams and crystals get smaller and smaller the use of multi-crystal data sets becomes unavoidable. The purpose of the challenge presented here was to represent a situation where the user decided to take relatively long exposures for each image in order to ensure that the high-resolution spots were visible to the eye. For small crystals, however, much of the sample’s useful life is used up in the first few images using this strategy (Evans *et al.*, 2011), and the challenge is to re-assemble all the data from a large number of highly incomplete collection runs, or wedges.

## 2. Methods

### 2.1 Preparation of simulated structure factors (F_right_)

Although it is possible to input F_obs_ into a MLFSOM (Holton *et al.*, 2014) simulation, F_obs_ is seldom 100% complete, and any missing hkls provided to MLFSOM will be taken as zero when rendering the simulated images and thus image processing software will assign them a well-measured intensity of zero. This will happen even if the reason for the missing F_obs_ was because the spot was saturating the detector in the original experiment, which is a very large and unnatural systematic error. In addition, the anomalous differences of F_obs_ are invariably noisy, and often unavailable. For these reasons, it is convenient to use calculated structure factors, which are always 100% complete, have a well-known phase and, by definition, no error in the amplitudes. Additional systematic errors can then be clearly defined and applied, depending on the goals of the simulation.

Calculated structure factors such as those output from refinement programs are typically denoted F_calc_, but for clarity here the symbol F_right_ shall denote the calculated structure factors that are fed into an image simulator. Thus F_right_ denotes the “right answer” used to evaluate data processing results. Structure factors obtained from simulated images shall be denoted F_sim_, as opposed to F_obs_, which will be reserved for actual real-word experimental observations. The distinction is important because the largest source of systematic error in macromolecular crystallography lies in the difference between F_obs_ and F_calc_, but the exact origin of this “R factor gap” remains unclear (Holton *et al.*, 2014). Specifically, refinement against F_right_ or F_sim_ derived from a simple single-conformer model invariably converges to abnormally low R_work_ and R_free_ after automated building and refinement. This is a glaring inconsistency with real data, and potentially makes the simulated data unrealistically easy to solve, diminishing its usefulness in benchmarking and debugging. More realistic R factors can be obtained by adding random numbers to F_right_, but the appropriate random distribution to use is not clear. Instead, values of F_right_ were generated here to have a combination of physically plausible systematic errors, and one final empirical systematic error.

### 2.2 I1 domain from titin (1g1c): lysozyme’s evil twin

Titin I1 domain was selected because its PDB entry 1g1c has unit cell, a=38.3 b=78.6 c=79.6 Å, which is the closest non-tetragonal unit cell to that of tetragonal *gallus gallus* egg lysozyme. The true space group is P2_1_2_1_2_1_, and thus represents an excellent challenge to software projects that seek to resolve indexing ambiguity in multicrystal projects, automatic space group assignment, and identification of crystallization contaminants by searching for similar unit cells in a database (McGill *et al.*, 2014, Simpkin *et al.*, 2018).

Coordinates and observed structure factor data for 1g1c were downloaded from the PDB (Berman *et al.*, 2002) and the cif-formatted structure factor data converted to mtz using the CCP4 Suite program CIF2MTZ (Winn, 2003). This mtz file header was edited with MTZUTILS to make a=38.3 and b=c=79.1 Å. The deposited coordinates were then refined against this new mtz file using phenix.refine (Adams *et al.*, 2010) for three macro-cycles.

This single-conformer model was used to compute F_right_ for a preliminary MLFSOM simulation, but downstream analysis suffered from the unrealistically low R_free_ < 2% statistics mentioned above. Previous studies (Holton *et al.*, 2014) found that using F_right_ from a multi-conformer model leads to more realistic R_free_, but modern building programs, such as qFit (van den Bedem *et al.*, 2009), can easily identify 2 or 3 alternate conformations. Real crystals contain trillions of different conformations, but approximating them as a Gaussian distribution simply recovers a canonical B factor.

Therefore, in order to create physically plausible systematic error that is not easily captured by automated building, 20 alternate conformations were generated for this simulation.

Twenty new pdb files were created from the single-conformer reference by perturbing each atom position, including all waters, with a random coordinate shift consistent with the assigned atomic B factor (B_atom_) using the jigglepdb.awk script distributed with MLFSOM (Holton *et al.*, 2014). Specifically, each coordinate shift along x,y and z was taken from a Gaussian distribution with root-mean-square (RMS) variation equal to sqrt(B_atom_/24)/π, which is the RMS shift that recapitulates the B factor at infinite trials. For example, consider a carbon atom at rest, which means B_atom_ = 0. A simple way to account for atomic motions would be to shift this atom about by say RMS 0.318 Å, and for each of thousands of positions and average the electron density. A less expensive way to get the same map, however, is to apply a B factor of 24. These perturbed coordinate sets therefore represent the natural deviations expected to be found from unit cell to unit cell in the crystal. Each of the 20 perturbed models was then refined against the re-indexed F_obs_ data using phenix.refine (Adams *et al.*, 2010) for ten macro-cycles with no free-R flags. This operation allowed the coordinates to relax away from any clashes and geometric distortions due to the unit cell change and random coordinate shifts and at the same time become more consistent with F_obs_. The reason for disabling free R flags was to avoid creating an artificial R_work_ vs R_free_ bias in F_right_. The final RMS deviation between these 20 re-refined models ranged from 0.75 to 0.9 Å (0.27 to 0.34 Å for Cα only). Each re-refined model was then edited to change all the methionine sulfur atoms to selenium. The refined solvent parameters: k_sol, B_sol, Rsolv and Rshrink were extracted from each phenix.refine run and then used with the selenium-containing coordinates in phenix.fmodel to generate 20 complete sets of calculated anomalous structure factors (F_model_) out to 1.8 Å resolution. These 20 F_model_ sets differed from each other by 14-20%, and were averaged together as complex numbers (F_avg_), and as root-mean-square amplitudes (F_rms_). The difference between F_avg_ and F_rms_ was that F_rms_ represented averaging in the intensity domain where the 20 F_model_ sets were assumed to come from different independently-diffracting mosaic domains. Conversely, F_avg_ was equivalent to averaging electron density maps, and assumed all 20 structures could be found within the coherence length of the beam. The R factor between F_avg_ and F_rms_ was only 3.3%, but since F_rms_ represents a physically plausible systematic error, it was carried on to the next step.

An empirical “R factor gap” systematic error was extracted by refining the deposited 1g1c model against the deposited 1g1c data and taking the F_obs_-F_calc_ amplitude difference for all observed reflections (F_diff_). F_diff_ was taken to be an empirical systematic error and added to F_rms_ to form F_sys_. Reflections missing F_obs_ were given F_diff_=0, and the resulting R factor between F_rms_ and F_sys_ was 18%. Finally, resolution was made to be slightly better than that available in 1g1c by applying a B factor of −15 to F_sys_ to form the value of F_right_ that was fed into the MLFSOM (Holton *et al.*, 2014) simulation.

### 2.3 Image simulation runs

Image simulations were conducted with MLFSOM (Holton *et al.*, 2014) using parameters matching the behavior of an Area Detector Systems (ADSC, Poway, CA) model Q315r x-ray detector, which is essentially a powdered Gd_2_O_2_S phosphor bonded to a charge-coupled device (CCD) via a fiber-optic taper (Holton *et al.*, 2012, Gruner *et al.*, 2002, Gruner, 1989, Waterman & Evans, 2010). These parameters were: electro-optical gain of 7.3 CCD electrons per X-ray photon, amplifier gain of 4 electrons per pixel intensity unit (ADU), zero-photon pixel level or “ADC offset” set to 40 ADU, and read-out noise of RMS 16.5 electrons/pixel. An intensity vignette falling to 40% at the edge of each module was used, and the Moffat function for the fiber-coupled CCD point-spread function as described in Holton *et al.* (2012) was varied from a *g*-value of 30 microns at the center of each module to 60 microns at the corner. Calibration error was set to RMS 3% with a spatial period of 50 pixels. This is in contrast to the true detector behavior of sub-pixel calibration error (Waterman & Evans, 2010), but was found in previous simulations to produce realistic R_merge_ values.

Image header values were made to be exact, with the exception of the beam center, which always requires further qualification. The header value was x,y = 154.96, 155.75 mm, which is one pixel off in each direction from the true beam center (155.063, 155.647) in the convention of the ADXV (Szebenyi *et al.*, 1997, Arvai, 2012) diffraction image viewer program. This 1-pixel shift is an example of the unfortunately common array of caveats that can enter into a beam center. Switching between programs that start counting pixels at 1 vs zero will generate 1-pixel shifts, and changing the definition of a pixel location from its center to one of the corners results in half-pixel shifts. More serious changes in beam center convention involve swapping the x and y axes, changing the origin among the four corners of the image, and two possible mirror flips. Different processing programs have different conventions, and despite significant efforts to standardize (Parkhurst *et al.*, 2014), do not always recognize and convert header values properly. Correct values were: x_beam 159.353, y_beam 155.063 for denzo/HKL2000 (Otwinowski & Minor, 1997), BEAM 159.301 155.011 for MOSFLM (Leslie & Powell, 2007), ORGX= 1512.73 ORGY= 1554.57 for XDS (Kabsch, 2010), and origin= −155.063, 159.356, −250 for CCTBX/DIALS (Grosse-Kunstleve *et al.*, 2002, Winter *et al.*, 2018). Note that in addition to the x-y flip between the ADXV and MOSFLM/HKL2000 convention there is a half-pixel difference between the conventions of MOSFLM and HKL2000, and a one-pixel difference between the MOSFLM and XDS conventions. Also, the XDS and DIALS conventions do not use the beam itself as a reference point, so the values provided above are appropriate only when other program settings declare the detector plane perfectly orthogonal to the incident beam. This is usually the case at the start of processing, but refinement of detector tilt will change these origin values. Detector tilts were simulated, but not included in the image header, specifically: 0.365708° forward detector tilt, 0.1145° detector twist, and −0.140959° of detector rotation about the beam (CCOMEGA) as defined in the MOSFLM convention (Leslie & Powell, 2007), and finally 0.0951363° rotation of the spindle about the vertical axis, away from normal to the beam. Although these numbers have many decimal places, they are the exact values fed into the simulation.

A total of 100 random orientation matrices with no orientation bias were pre-generated and used to create 100 simulated runs of 15 images each. Each run, or “wedge”, began with a new, fresh crystal that was assigned a cube shape with edge dimension selected randomly about a 5 micron average value and RMS 1 micron variation. Crystals larger than 6 microns were cut off by the 6 micron wide square beam, and this illuminated volume did not change with rotation. Final illuminated volumes are listed in Table 1. The X-ray beam was made to have flux 1×10^12^ photons/s into a 6 micron wide flat-top profile. Per-image exposure time was 1 s and ΔΦ = 1°. Shutter jitter was set to RMS 2×10^−3^ s in the starting and ending Φ values of each image while beam flicker was taken to be 0.15%/√Hz and implemented in 10 steps per second. Beam divergence was set to 0.115° x 0.0172° (horiz x vert). These are typical measured properties of the Advanced Light Source beamline 8.3.1 (MacDowell *et al.*, 2004). Spectral dispersion, however, was set to 0.3% instead of the 0.014% measured form the Si(111) monochromator in order to mimic unit cell variations in the sample (Nave, 1998). The mosaic spread was set to be a uniform disk of sub-crystal orientations with diameter 0.23°.

**Table 1:** Simulated crystal volumes (µm^3^) The true scale factor of the spots from each simulated dataset is directly proportional to the simulated crystal volume, which was chosen randomly for each crystal. The actual values used in the simulation are listed here and may be used to check the accuracy of scaling programs because no other variables such as x-ray beam flux, or even the structure factors were varied from crystal to crystal. The only remaining correction after this is the resolution dependent scale factor of the simulated radiation damage described in §2.4.

| xtal | volume | xtal | volume | xtal | volume | xtal | volume | xtal | volume |
| --- | --- | --- | --- | --- | --- | --- | --- | --- | --- |
| 001 | 225 | 021 | 139 | 041 | 132 | 061 | 50.3 | 081 | 105 |
| 002 | 56.3 | 022 | 232 | 042 | 234 | 062 | 99.5 | 082 | 230 |
| 003 | 63.9 | 023 | 155 | 043 | 46.9 | 063 | 196 | 083 | 171 |
| 004 | 220 | 024 | 114 | 044 | 75.9 | 064 | 102 | 084 | 122 |
| 005 | 186 | 025 | 38.4 | 045 | 51.6 | 065 | 229 | 085 | 56.8 |
| 006 | 89.2 | 026 | 155 | 046 | 89.1 | 066 | 161 | 086 | 90.5 |
| 007 | 52.2 | 027 | 46.7 | 047 | 230 | 067 | 72.4 | 087 | 90.2 |
| 008 | 249 | 028 | 60.7 | 048 | 56.7 | 068 | 14.5 | 088 | 171 |
| 009 | 185 | 029 | 70.7 | 049 | 97.8 | 069 | 131 | 089 | 186 |
| 010 | 110 | 030 | 166 | 050 | 153 | 070 | 37.5 | 090 | 128 |
| 011 | 166 | 031 | 143 | 051 | 237 | 071 | 207 | 091 | 42.2 |
| 012 | 121 | 032 | 132 | 052 | 87.4 | 072 | 159 | 092 | 295 |
| 013 | 160 | 033 | 213 | 053 | 130 | 073 | 88.4 | 093 | 240 |
| 014 | 60.4 | 034 | 27.8 | 054 | 128 | 074 | 60.2 | 094 | 148 |
| 015 | 189 | 035 | 210 | 055 | 86.4 | 075 | 190 | 095 | 51.5 |
| 016 | 39.4 | 036 | 100 | 056 | 127 | 076 | 39.2 | 096 | 134 |
| 017 | 47.6 | 037 | 12.5 | 057 | 52.8 | 077 | 186 | 097 | 46.3 |
| 018 | 123 | 038 | 228 | 058 | 104 | 078 | 78.5 | 098 | 15.8 |
| 019 | 277 | 039 | 210 | 059 | 146 | 079 | 108 | 099 | 201 |
| 020 | 71.4 | 040 | 83.7 | 060 | 102 | 080 | 31.2 | 100 | 111 |

The X-ray background was also rendered on an absolute scale using realistic thicknesses of materials in the beam: 20 mm of helium gas between the collimator and beam stop, 5 microns of liquid water, and 4 microns of Paratone-N oil in the beam path. Compton and diffuse scatter from the crystal lattice itself were computed based on the size and composition of the macromolecule as described in the supplementary materials of Holton *et al.* (2014). Briefly, at the resolution where the Bragg spots fade into the background this diffuse component of the background converges to the same level expected from all the atoms in the protein crystal scattering independently, as if they were a gas.

### 2.4 Simulated Radiation Damage Model

Radiation damage was simulated in MLFSOM (Holton *et al.*, 2014) using only a simple, resolution-dependent exponential decay of spot intensities with dose using Equation 13 from Holton and Frankel (2010):

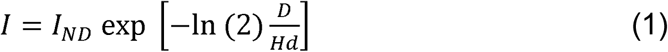

where: *I*_*ND*_ is the intensity that would be observed in the absence of radiation damage, *I* is the spot intensity at dose *D* (MGy), *d* is the resolution of the spot (Å), and *H* is the 10 MGy/Å estimated by Howells *et al.* (2009). For example, spots in the simulation at 2 Å resolution were made to fade exponentially with dose reaching half of *I*_*ND*_ after 20 MGy, and spots at 3.5 Å faded by half at 35 MGy. Dose was calculated assuming the crystal was bathed in a flat-top beam using the formula 2000 photons/um^2^/MGy from Holton (2009). This puts the first image at 13.9 MGy (see Fig 1), and it should be noted that this end-of-image dose was used for the average dose of the entire image. No attempt was made to average over sub-image decay for this simulation, and the result was that the decay curve appears to be a perfect exponential offset in dose by half an image. Non-isomorphism due to radiation damage was not simulated, and except for the simple exponential spot-fading described above no variation in structure factors or unit cell with dose was employed. In fact, the unit cell and structure factor table was identical for all 100 simulated crystals, making this a case of perfect isomorphism. The reason for these unrealistically perfect damage and isomorphism models was to simplify estimation of the errors in the cell and damage model introduced by the simulated noise as well as the data processing algorithms themselves.

It is noteworthy that although (1) is consistent with 13 distinct studies of crystals and single particles using both x-rays and electrons surveyed by Howells *et al.* (2009) over a resolution range of 2 Å to 600 Å, it is not equivalent to a B factor that increases with dose. This is incongruous with popular scaling programs, which use a quadratic (B factor) rather than linear (1) resolution dependence to spot fading (Blake & Phillips, 1962, Evans, 2006). This damage model is therefore an example of a systematic error between the simulation and the internal models of scaling programs. The nature of the systematic error between these models and reality is no doubt more complex, but here we simply used the average trend of spot fading vs resolution as the sole manifestation of radiation damage.

**Figure 1:**
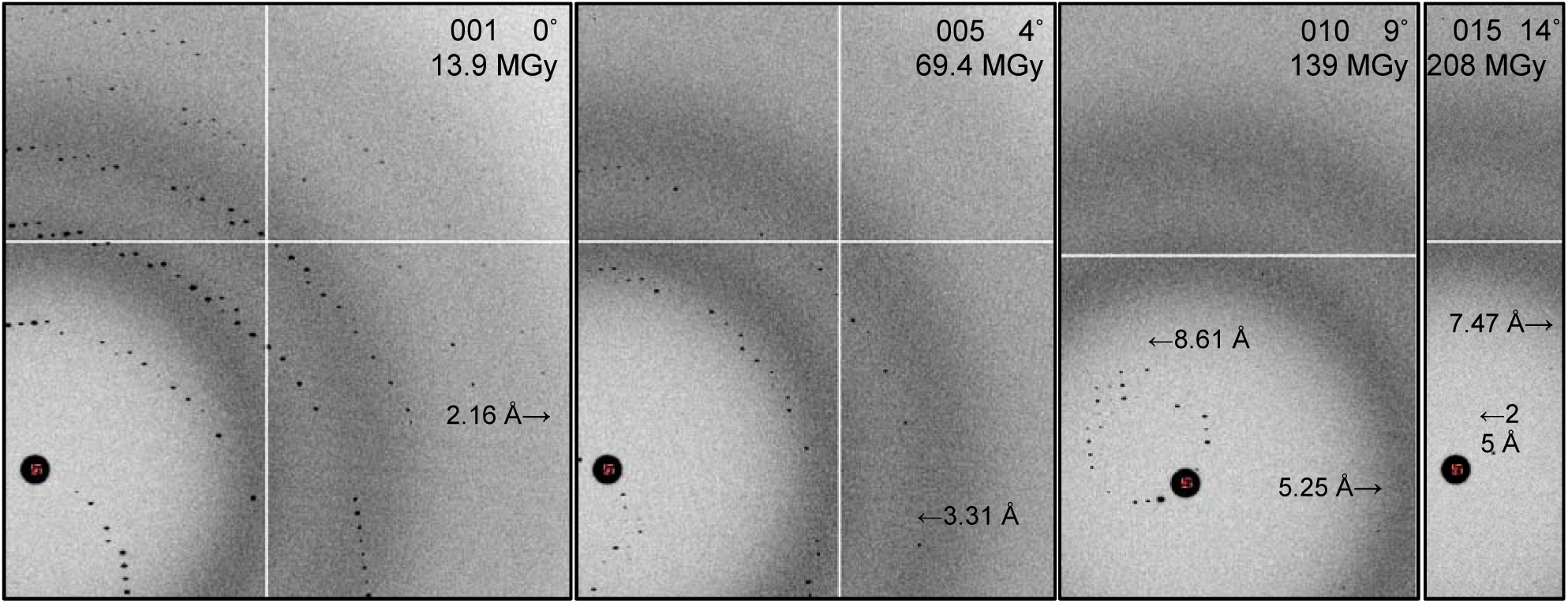
Zoomed-in sections of diffraction patterns from simulated crystal 016. Six lunes are apparent on image 001, but indexing this wedge still proved problematic. The resolution dependent exponential fading of spots with is exemplified by the rapid loss of high-angle data and relative persistence of low-angle features. Despite perfect isomorphism, images 004 and higher degraded overall anomalous signal, and images 002 and higher degraded the overall resolution of the final data set.

## 3. Results and Discussion

In order to demonstrate the utility of this challenge some discussion of the difficulties encountered when trying to process the images using MOSFLM (Leslie & Powell, 2007), LABELIT (Sauter & Poon, 2010), HKL2000 (Otwinowski & Minor, 1997), XDS/XSCALE (Kabsch, 2010), DIALS (Winter *et al.*, 2018), Phenix (Adams *et al.*, 2010) and the CCP4 Suite (Winn, 2003) are described here. Specific bugs and program to program differences will not be detailed since these programs are continuously improving and contemporary shortcomings have little archival value, but the algorithmic challenge of simultaneous speed and robustness will be evaluated.

### 3.1 Automatic indexing

Despite the high degree of similarity between these 100 simulated crystals automated indexing was not always successful. Depending on the software used, choice of images, and settings for spot picking and cell restraints, failures ranged from exiting with an error message to confidently arriving at an incorrect Niggli cell, usually with one or more of the primitive cell dimensions doubled. This type of mis-indexing could not be corrected by downstream re-indexing programs such as POINTLESS (Evans, 2006, 2011), and thus represents a significant barrier to including these particular wedges.

A naive user might even mistake such mis-indexing for evidence of variations in crystal habit, so it is important to note here that there was no difference in quality between any of these simulated crystals. All wedges had the same resolution, same decay rate, and were perfectly isomorphous. Illuminated volume varied over a factor of 24, but neither the smallest (037) nor largest (092) simulated crystal had indexing problems. The most problematic crystals were 016, 064, 065, 086, and 095, all of which have one reciprocal cell axis close to parallel with the incident beam. This situation can cause problems in indexing because the information about the cell axis near the beam is maximally distorted by the Ewald sphere, and may even be missing entirely if the crystal diffracts poorly and produces only one lune (Brewster *et al.*, 2018). However, all these problematic wedges diffracted to 1.8 Å resolution and displayed 3-6 clear lunes, so the reason for these failures is not immediately clear. In addition to these five problem crystals, four others: 051, 054, 062, and 063 failed with most combinations of images but not all, and 11 more: 004, 006, 010, 019, 065, 068, 086, 094, 097, and 098 would usually succeed, but failed with at least one combination of images. Since the major difference was the crystal orientation, the indexing algorithm itself may be considered a source of orientational bias in multi-crystal data, even if the true orientation distribution is isotropic.

In general the fastest programs had the highest failure rates, whereas more complex algorithms would take longer but arrive at the correct Niggli cell more reliably, such as that of Sauter and Zwart (2009). Execution times varied from 0.3 s to 9 s across the programs tested, so the tradeoff between speed and robustness is significant. However, these same more complex algorithms were vulnerable to other considerations, such as weak images. For example, LABELIT indexing with images 1 and 15 reproducibly failed, but the same program given images 1 and 4 would succeed. A combinatorial approach scanning over image selection and other program settings would no doubt be most robust, but also consume the most computing resources.

Automatic space group determination also had its flaws. Essentially all indexing software tested arrived at a tetragonal solution, which is not intrinsically problematic until after the merging step, but the completeness of any given single wedge was so low ∼10% that few symmetry operators can be eliminated for any particular wedge taken in isolation. For example, POINTLESS (Evans, 2006, 2011) assigned most of the 100 simulated crystals to space groups P1 (35%) or P2 (23%), some were assigned to P222 (11%), C2 (12%), or P422 (9%) and in rare cases C222 or P4, indicating that the true space group is not obvious from the primary data. It is commonplace to assign the highest symmetry possible during processing in order to maximize the completeness of each wedge and therefore the overlap with other wedges to make cross-crystal scaling simpler and more robust. However, pursuing this strategy invariably ended with what appeared to be extremely noisy data that did not merge well and appeared to be twinned. Final R factor between F_sim_ and F_right_ was 53%. The most robust strategy and unfortunately the most computationally intensive remained independently pursuing processing, scaling, merging and combining data in all possible point groups separately, and in addition scanning over all possible radiation damage cutoffs. This is a large number of combinations, but the correct point group (222) and cutoff (3 images) were only clear when both were applied at the same time.

### 3.2 Cheating

In order to demonstrate an ideal solution to this challenge the simulated data were processed using F_right_ as a reference for the unit cell and structure factors. This eliminated any indexing ambiguity. The unit cell and space group were also fixed to the correct values during indexing, refinement and integration in MOSFLM. The best radiation damage cutoff was determined empirically by scaling and merging all 100 correctly-indexed wedges together with POINTLESS/AIMLESS (Evans, 2011), and comparing the final merged structure factors to F_right_. The optimum cutoff to optimize weak, high-resolution data was to use only the first image, as evidenced in Fig 2. Although scaling programs such as AIMLESS are designed to take a “run” of images, for this case each run started and ended with image “1”, a strategy that also eliminates all partially-recorded reflections. Using just the first image from each wedge also minimized the overall R_work_ = 21.3% and R_free_ = 25.7% after refining the selenated 1g1c reference model to convergence with REFMAC (Murshudov *et al.*, 2011).

**Figure 2:**
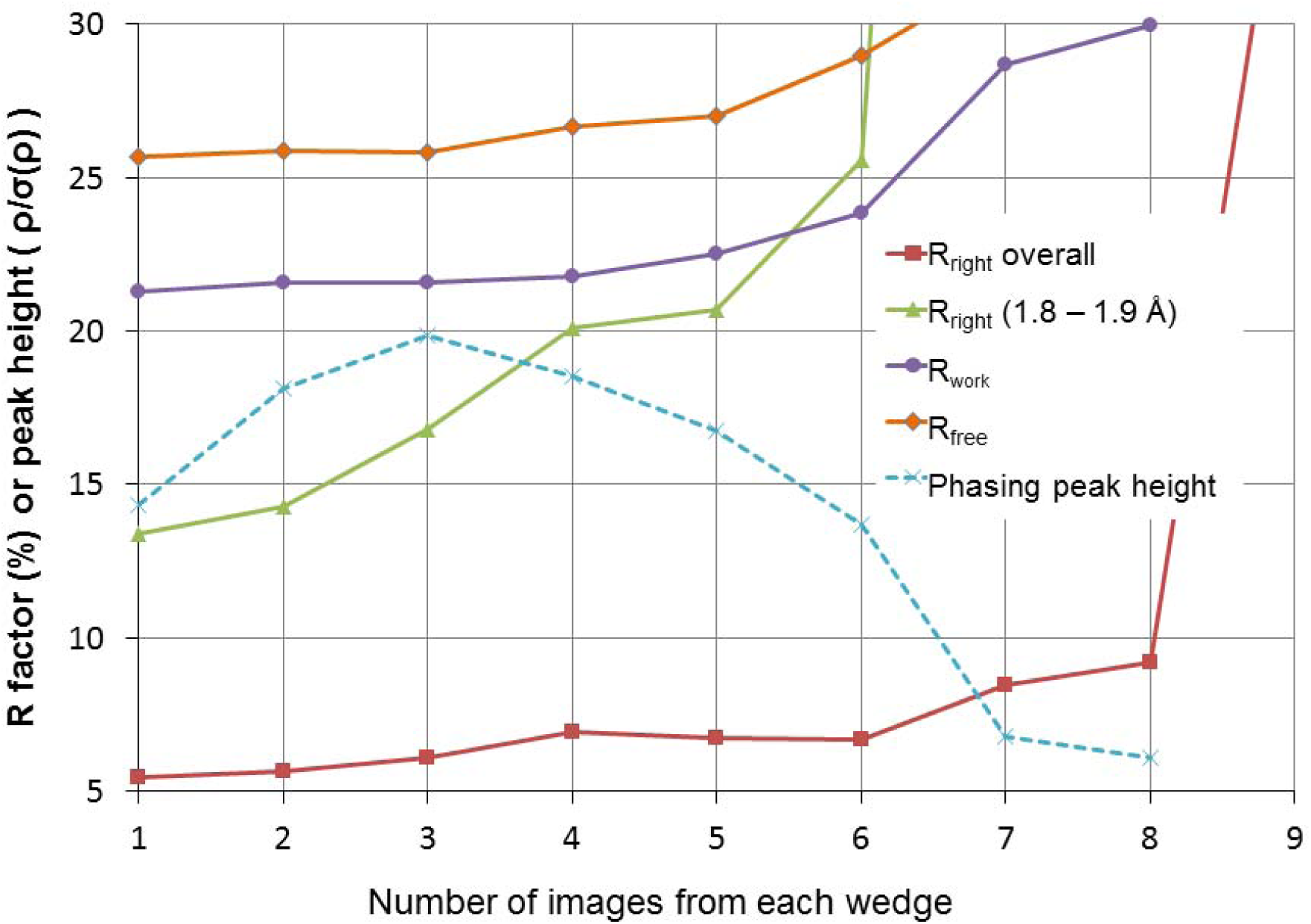
Graph of relative error (R_right_) between correct structure factor (F_right_) and the structure factor obtained from scaling and merging the first N images from all 100 simulated crystals (F_sim_). N is the x-axis. Also shown are R_work_ and R_free_ from refinement to convergence of the correct starting model against F_sim_ from N-image data. Despite perfect isomorphism, fewer images resulted in better agreement. The y axis also represents the maximum peak height found in the phased anomalous difference Fourier (dashed line). Phases were obtained by removing all selenium atoms before refining to convergence against F_sim_. The phasing signal is maximized at N=3.

The optimum anomalous signal was attained using the first 3 images of each wedge (Fig 2), and automated structure solution was straightforward using the SHELXC/D/E pipeline (Sheldrick, 2008). Structure solution was also still possible using crystals 001-071, but not 001-070, indicating the threshold of solvability with ideal data processing. It may be possible to apply further cheats, such as directly correcting the exponential spot decay and get better results, but this was not attempted in the present work.

### 3.3 Resolution dependence of radiation damage

The non-Gaussian nature of the damage model used in this simulation was unexpectedly detrimental to contemporary scaling procedures, so here we shall place this empirical decay equation into context with the conventional scale-and-B-factor model. It is instructive to re-cast (1) in the same form as a B factor (exp(-*Bs*^2^)) by defining *A* = ln(2)*D/H*, substituting the resolution *d* with the reciprocal scattering vector length *s* = (2*d*)^−1^, and converting intensities (*I*) to structure factors (*F*) by taking the square root of both sides. The factor of two in the switch from *d* to *s* is cancelled by the switch from intensities to structure factors, and we arrive at:

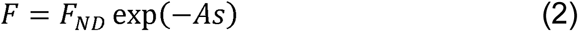

where: *F*_*ND*_ is the structure factor of the damage-free unit cell. This “A factor” was first described in a lecture by this author at the 7^th^ International Workshop on Radiation Damage to Biological Crystals and was subsequently published by Borek *et al.* (2013). This re-arranged spot-fading formula immediately suggests a Taylor expansion in the exponent demonstrating the relationship between *A* and *B* and perhaps additional factors, such as *C*. Let us briefly entertain this formalism, and write:

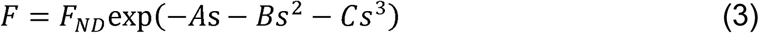

where: *B* is the usual B factor 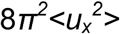 with *u*_*x*_ as the component of the Gaussian-distributed atomic displacement vector ***u*** in the direction normal to the Bragg plane and <> denotes the mean over all atoms. Similarly, *A* = 2π*w*_*fhm*_ where *w*_*fhm*_ is the full-width at half-maximum of atomic displacements taken from the multivariate Cauchy-Lorentz distribution:

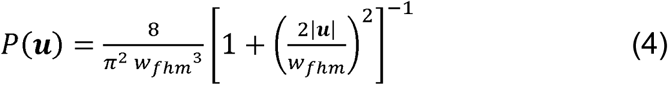

where: *P*(***u***) is the normalized probability of atomic displacement vector ***u***, and || denotes vector magnitude (in Å). This distribution resembles a Gaussian but has heavier tails, indicating a much higher ratio of large-scale to small-scale movements than would be expected from a Gaussian distribution. Generating this distribution must be done with care because one cannot simply apply three independent displacements along x,y and z, as this creates a highly anisotropic 3D histogram. Rather, a random direction for ***u*** must first be chosen and (4) applied along its axis.

It was argued by Debye (1914) that all terms except *Bs*^*2*^ in (3) vanish when averaged over the large number of atoms in the crystal (James, 1962 Eq. I.26), but this is only the case when the distribution of atomic displacements converges to a Gaussian via the Central Limit Theorem. There are random distributions that do not obey the Central Limit Theorem, and the Cauchy-Lorentz is one example. In fact, combinations of Cauchy-Lorentz deviates always converge to another Cauchy-Lorentz distribution, forming an analogous but distinct version of the Central Limit Theorem.

Strictly speaking, the fall-off of intensity with resolution due to any distribution of atomic displacements is the Fourier transform of that distribution. The Fourier transform of a Gaussian atomic displacement distribution is another Gaussian (the *B* factor), and the Fourier transform of a Cauchy-Lorentz distribution is an exponential in reciprocal space, as in (2). If the manifestation of radiation damage is a B factor that increases linearly with dose then the spot-fading half-dose should be related to the square of resolution, not linearly. The Howells observation of a linear relationship between resolution and spot-fading half-dose therefore implies a direct proportionality between dose and the width of the distribution of atomic displacements:

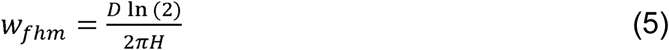

where: D is the dose in MGy, ln(2) is the natural log of 2 and H is the 10 MGy/Å trend observed by Howells. Here we use the full width at half max to describe the Cauchy-Lorentz histogram rather than the RMS variation because the RMS variation of a Cauchy-Lorentz distribution is undefined, as is it’s mean. A physically reasonable explanation for the departure from Gaussian-distributed atomic displacements may be that large enough displacements require neighboring atoms to move out of the way, creating additional large ***u*** vectors of similar magnitude and direction and leading to a higher than “normally” expected population of large ***u*** vectors. Cracking and slipping of lattice fragments relative to each other may be examples of such concerted movements.

The appearance of the letter *B* as the second term in (3) suggests an origin for the choice of the letter *B* to indicate the Debye-Waller-Ott factor, and therefore a natural place for *A*- and *C*-factors. However, the *C*-factor does not correspond to any physically reasonable distribution because its corresponding real-space displacement histogram has negative population values, and probabilities cannot be negative. So, although (3) resembles a Taylor expansion in the exponent only the first two terms *A* and *B* correspond to physically plausible distributions. In addition, the first use of *B* to describe Debye’s disorder parameter appeared in Bragg (1914), and therein the letter *A* was used to encapsulate the overall scale factor, which is in no way analogous to the Cauchy-Lorentz term in (3). It is therefore recommended that the use of “A factor” as in Borek *et al.* (2013) be avoided, and instead replaced with the more informative expression: A = 2π*w*_*fhm*_.

## 4. Conclusions

The challenges to macromolecular structure determination using data from a large number of small crystals lie primarily in the combinatorial nature of the data analysis. Recent landmark achievements such as (Brehm & Diederichs, 2014, Liu & Spence, 2014, Gildea & Winter, 2018, Diederichs, 2017) represent important mathematical advances in handing this problem, but even these approaches are sensitive to incorrect indexing and radiation damage cutoffs during processing, leaving these as choices that must still be exhaustively pursued. By “cheating” this work was able to solve the challenge structure using only the first 71 crystals of the 100 presented, and further work that can lower this number without cheating will directly translate to real-world projects finishing earlier and using fewer difficult-to-produce isomorphous crystalline samples.

It is tempting to suggest overcoming indexing problems by using a pair of orthogonal alignment shots prior to data collection, but since only the first three images appear to be useful before data quality degrades this strategy is not recommended. Lowering the exposure time and gathering more of reciprocal space with the same dose is expected to improve indexing performance, but this strategy is not applicable to the problem of serial crystallography (Wiedorn *et al.*, 2018, Chapman *et al.*, 2011), where particularly at XFEL sources only one image is available from every sample. The limit of how weak individual images can be before resolution begins to degrade will be the subject of a future challenge but recent results have shown that this limit can be quite low (Lan *et al.*, 2018, Parkhurst *et al.*, 2016). It is further expected that as radiation damage processes become better understood and correctable including more images will improve data quality rather than degrade it.

## Acknowledgements

I would like to thank Drs. Christine Gee and Nicholas Sauter for extremely helpful discussions of this manuscript. This work was supported by grants from the National Institutes of Health (GM124149, GM124169 and GM103393) and the US Department of Energy under contract No. DE-AC02-05CH11231 at Lawrence Berkeley National Laboratory. Images have been deposited into the IRRMC at proteindiffraction.org and are also available at: http://bl831.als.lbl.gov/∼jamesh/challenge/microfocus/.

